# Deconvolution of multiple infections in *Plasmodium falciparum* from high throughput sequencing data

**DOI:** 10.1101/099499

**Authors:** Sha Joe Zhu, Jacob Almagro-Garcia, Gil McVean

**Affiliations:** Wellcome Trust Centre for Human Genetics, University of Oxford, Oxford, UK; Medical Research Council (MRC) Centre for Genomics and Global Health, University of Oxford, Oxford, UK; Wellcome Trust Sanger Institute, Hinxton, UK; Big Data Institute, Li Ka Shing Centre for Health Information and Discovery, University of Oxford, Oxford, UK

## Abstract

**Motivation:** The presence of multiple infecting strains of the malarial parasite *Plasmodium falciparum* affects key phenotypic traits, including drug resistance and risk of severe disease. Advances in protocols and sequencing technology have made it possible to obtain high-coverage genome-wide sequencing data from blood samples and blood spots taken in the field. However, analysing and interpreting such data is challenging because of the high rate of multiple infections present.

**Results:** We have developed a statistical method and implementation for deconvolving multiple genome sequences present in an individual with mixed infections. The software package *DEploid* uses haplotype structure within a reference panel of clonal isolates as a prior for haplotypes present in a given sample. It estimates the number of strains, their relative proportions and the haplotypes presented in a sample, allowing researchers to study multiple infection in malaria with an unprecedented level of detail.

**Availability and implementation:** The open source implementation *DEploid* is freely available at https://github.com/mcveanlab/DEploid under the conditions of the GPLv3 license. An R version is available at https://github.com/mcveanlab/DEploid-r.

## 1 Introduction

Malaria remains one of the top global health problems. The majority of malaria related deaths are caused by the *Plasmodium falciparum* parasite (WHO, 2016), transmitted by mosquitoes of the genus *Anopheles*. Patients are often infected with more than one distinct parasite strain (termed mixed infection, multiple infection, or complexity of infection), due to bites from multiple mosquitoes, mosquitoes carrying multiple genetic types or a combination of both. Mixed infections can lead to competition among co-existing strains and may influence disease development (de Roode et al., 2005), transmission rates (Arnot, 1998) and the spread of drug resistance (de Roode et al., 2004). In addition, within-host evolution can lead to the presence of more than one genetically and phenotypically distinct strains (Bell et al., 2006).

The presence of multiple strains of *P. falciparum* makes fine scale analysis of genetic variation challenging, since genetic differences between strains of this haploid organism will appear as heterozygous loci. Such mixed calls confound methods that exploit haplotype data to detect, among other phenomena, the occurrence of natural selection or recent demographic events (Harris and Nielsen, 2013;Lawson et al., 2012;Mathieson and McVean, 2014; Sabetil et al., 2002). In light of these difficulties, researchers usually focus on clonal infections or resort to heuristic methods for resolving heterozygous genotypes. The former approach discards valuable information regarding genetic diversity and relatedness, whereas the latter tends to create chimeric haplotypes that are not suitable for analysis, unless mixed calls are very sparse.

In comparison to the problem of phasing haplotypes within diploid organisms, deconvolving the strains of a multiple infection differs because of uncertainty in the number of strains present and their relative proportions. Consequently, existing tools for phasing diploid organisms, such as BEAGLE (Browning and Browning, 2007), IMPUTE2 (Howie et al., 2009) and SHAPEIT (Delaneau et al., 2012; O’Connell et al., 2016), are not appropriate. Galinsky et al. (2015) and O’Brien et al. (2015) have attempted to address the multiple infection problem by inferring the number and proportions of strains from allele frequencies within samples. However, since they do not infer haplotypes, these approaches have limited applicability.

As part of the Pf3k project (Pf3k, 2016), an effort to map the genetic diversity of *P. falciparum* at global scale, we have developed algorithms and a software package implementation `DEploid`, for deconvolving multiple infections. The program estimates the number of different genetic types present in the isolate, the proportion or abundance of each strain and their sequences (i.e. haplotypes). To our knowledge, `DEploid` is the first package able to deconvolute strain haplotypes and provides a unique opportunity for researchers to study the epidemiology of *P. falciparum.*

## 2 Methods

### 2.1 Notations

We first introduce our notation (see Table 1). Our data, *D*, are the allele read counts of sample *j* at a given site *i*, denoted as *r_j,i_* and *a_j,i_* for reference (REF) and alternative (ALT) alleles respectively. These are assigned values of 0 and 1 resepctively. Here we consider only biallelic loci, though future extension to include multi-allelic sites is simple. The empirical allele frequencies within a sample (WSAF) *p_j,i_* and at population level (PLAF) *f_i_* are calculated by 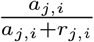 and 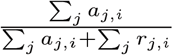 respectively. Since all data in this section refers to the same sample, we drop the subscript *j* from now on.

**Table 1.** Table summarising the notation used in this article.

|  |  |
| --- | --- |
| $i$ | Marker index |
| $j$ | Sample index |
| $r$ | Read count for reference allele |
| $a$ | Read count for alternative allele |
| $f$ | Population level allele frequency (PLAF) |
| $n$ | Number of strains within sample |
| $l$ | Sequence length |
| $\mathbf{w}$ | Proportions of strains |
| $\mathbf{x}$ | Log titre of strains |
| $\mathbf{h}_i$ | Allelic states of $n$ parasite strains at site $i$ |
| $h_{k,i}$ | Allelic state of parasite strain $k$ at site $i$ |
| $p$ | Observed within sample allele frequency (WSAF) |
| $q$ | Unadjusted expected WSAF |
| $\pi$ | Adjusted expected WSAF |
| $\Xi$ | Reference panel |
| $\xi_{k,i}$ | Allelic state of reference panel strain $k$ at site $i$ |
| $G$ | Scaling factor used for genetic map |
| $e$ | Probability of read error |

### 2.2 Model

We describe the mixed infection problem by considering the number of strains, *n*, the relative abundance of each strain, w, and their allelic states, h. Similar to O’Brien et al. (2015), we use a Bayesian approach and define the posterior probabilities of *n*, w and h given a reference panel, Ξ, and the read error rate, *e*, as:

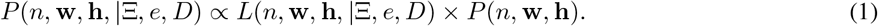

We assume a prior in which the haplotypes of the n strains are independent of each other and dependent only on the reference panel. Therefore, the joint prior can be written as:

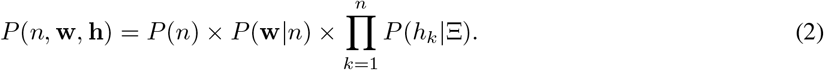

The following sections describe details of the model and the approach to inference.

#### 2.2.1 Likelihood function

Let w = [*w_1_,…, w_n_*] and h_*i*_ = [*h*_1,*i*_,…, *h_n_*_,*i*_] denote the proportions and alleic states of the *n* parasite strains at site *i.* We use O’Brien et al. (2015)’s expression for the expected WSAF at site *i, q_i_,* as:

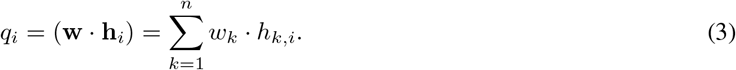

The data, which can be summarised by the reference and alternative allele read counts at each site, is modelled through a beta-binomial distribution given the expected WSAF. We model the data at distinct segregating sites as independent. Thus the likelihood function in Eqn. (1) is only dependent on the haplotypes present and their frequencies through their contribution to *q_i_*.

To incorporate sequencing error, we modify the expected WSAF such that the allele frequency of ‘REF’ read as ‘ALT’ is (1−*q_i_*)*e*, and the allele frequency of ‘ALT’ read as ‘REF’ is *q_i_e*. Thus, the adjusted expected WSAF becomes:

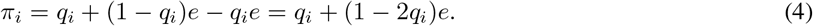

We model over-dispersion in read counts relative to the Binomial using a Beta-binomial distribution. Specifically, the read counts of ‘ALT’ are identically and independently distributed (i.i.d.) Bernoulli random variables with probability of success *v_i_*; i.e. *a_i_* ~ *Binom*(*a_i_* + *r_i_, v_i_*), and *v_i_* ~ *Beta*(*α, β*), where *E*(*v_i_*) = *α*/(*α* + *β)* = *π_i_*. This is achieved by setting *α* = *c* · *π_i_* and *β* = *c* · (1 − *π_i_*), such that the variance of the WSAF is inversley proportion to *c*. Combined, we have:

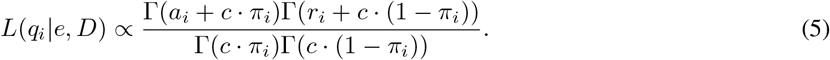

#### 2.2.2 Prior distributions

Rather than model the number of strains, *n*, directly, we take the approach of fixing *n* to be at the upper end of what can realistically be inferred (typically 5), using a skewed prior for proportions (such that typically only 1 - 2 strains might be at appreciable frequency) and then discarding strains inferred to have a proportion less than some critical amount (e.g. 1 percent).

To achieve this, we model the proportions of the *n* strains through a log titre, *x_k_*, drawn from a *N*(*η, σ*^2^) prior. The proportion of strain *k*, *w_k_*, is given by

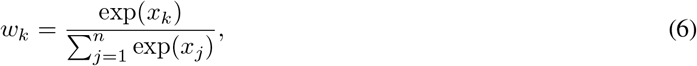

and the prior density is given by the distribution function for the value of x.

Haplotypes, h, are modelled as being generated independently from the reference panel by the Li and Stephens (2003) process, though with a rate of mis-copying that is independent of the panel size. That is, under the prior, a path through the reference panel is sampled as a Markov process where recombination enables switching between members of the reference panel and mis-copying allows the allelic state of the haplotype within the sample to differ from the allelic state of the reference panel haplotype being copied at the site. The transition probability of switching from copying reference haplotype *a* to reference haplotype *b* is (1 − exp(−*Gψi*))/|Ξ|, where *ψ_i_* is the genetic distance (in Morgans) between sites *i* and *i* + 1, *G*, is a scaling factor (described below in more detail) and |Ξ| is the size of the reference panel. Note that unlike the original model, the recombination or switching rate is not dependent on sample size.

For miscopying, let ξ_*k*_ denote the state of the sequence in the reference panel Ξ that *h_k_* is copying from at given site and *μ* denote the probability of miss-copying:

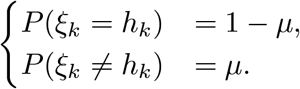

As above, this is a simple reparamterisation of the original model, but where the miscopying rate is independent of the sample size. The emission probabilities are given by the convolution of the reference panel paths and the miscopying process, strain proportions and the read error rate.

### 2.3 Inference

To perform inference about the haplotypes present and their proportions we use Markov chain Monte Carlo (MCMC). We use a Metropolis-Hastings algorithm to sample proportions (w) given h; and use a Gibbs sampler to update h for a given w, with two types of update: a single haplotype and a pair of haplotypes.

#### 2.3.1 Metropolis-Hastings update for proportions

We update w|*n*, through the underlying log titres, x|*n*. Specifically, we choose *i* uniformly from *n* and propose new 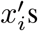 from 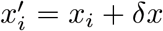 where *δx* ~ *N*(0, *σ*^2^/*s*), and *s* is a scaling factor. The new proposed proportion is therefore 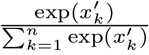. Since the proposal distribution is symmetrical, the Hastings ratio is 1. A new update is accepted with probability

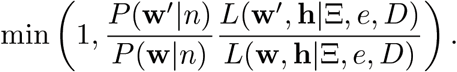

#### 2.3.2 Gibbs update for single haplotype

We choose haplotype strain *s* uniformly at random from *n* strains to update. At each site, given the current proportions, we can calculate the likelihood of the 0 and 1 states. To achieve this, we first remove it from the current WSAF, i.e. subtract *w_s_* · *h_s_* from Eqn. (3), which gives

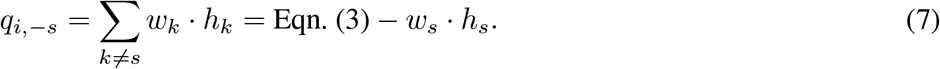

Therefore, updating the allelic state of strain *s* to 0 and 1, the expected WSAF becomes

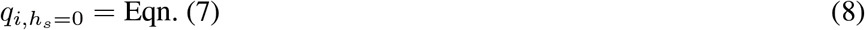

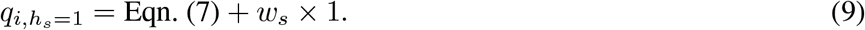

We substitute Equations (8) and (9) into Equation (5) after adjustment for read error.

Given the structure of the hidden Markov model and the above likelihoods, the forward algorithm can be used to sample a path through the reference panel, and subsequent mis-copying, efficiently from the marginal posterior distribution. In effect, the reference panel is used as a prior on haplotypes present in the sample (with recombination creating a mosaic of the different haplotypes) and the mis-copying process allows for recent mutation, recurrent mutation, gene conversion and some types of technical error. Figure 1 illustrates the approach.

**Figure 1.**
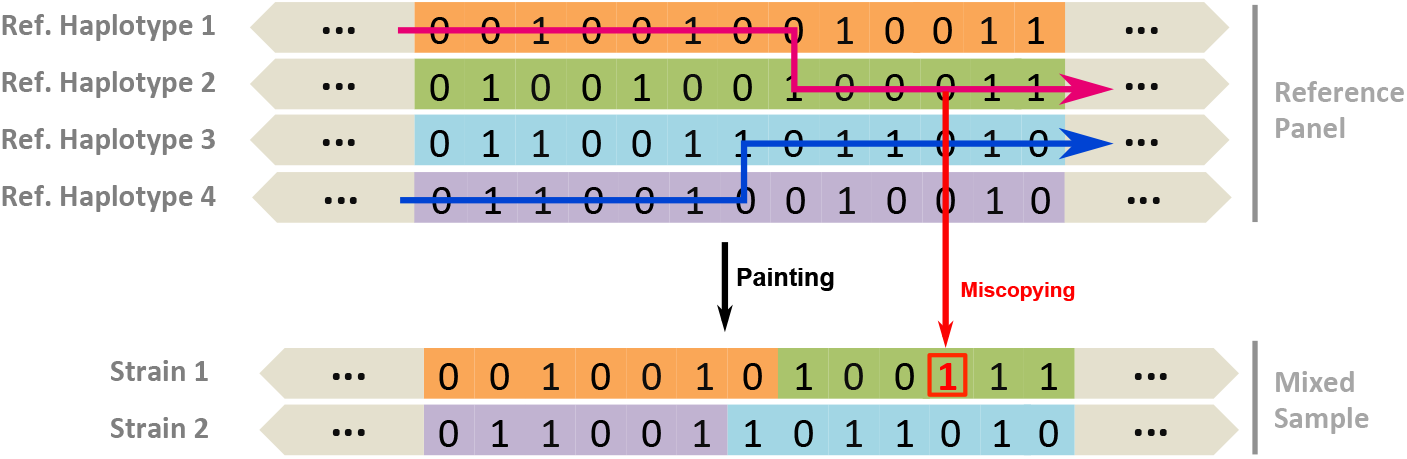
The Li and Stephens (2003) algorithm as applied to the problem of multiple strain inference. Strain 1 haplotype is made up from reference haplotype segments of 1 and 2; and strain 2 haplotype is made up from reference haplotype segments of 3 and 4. With mis-copying, we allow strain states differ from the path: At the third last position of strain 1, the path is copied from reference haplotype 2, with the state of 0.

#### 2.3.3 Gibbs update for a pair of haplotypes

In order to improve mixing, we also perform Gibbs-sampling updates for pairs of haplotypes (given current proportions). The algorithm proceeds as for the single-haplotype update, though with a larger state space. First, we sample a pair of haplotypes, *s*_1_ and *s*_2_, uniformly. As in Equation (7), we first remove their states from the WSAF:

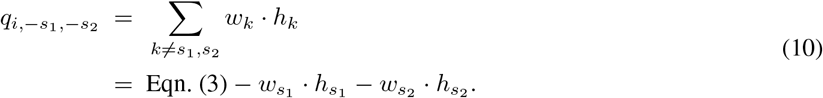

Considering all four possible combination of genotypes, we can then write down the expected WSAF:

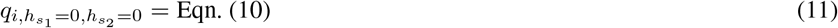

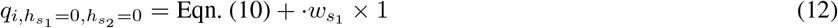

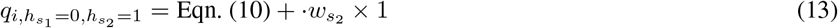

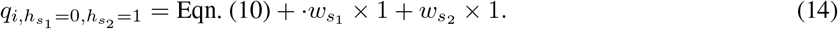

Substituting expressions. (11) to (14), into Equation (5), we then obtain their associated likelihoods.

As in the single-haplotype update, the hidden Markov model formulation enables us to sample a pair of paths through the reference panel (and the mis-copying process) efficiently from the marginal posterior distribution using the forward algorithm, that is given the other haplotypes and their inferred proportions. Equations describing the calculations are given in the Supplementary Material.

### 2.4 Implementation details

- **Number of strains.** As described above, we aim to infer more strains than are actually present, starting the MCMC chain with a fixed *n*, which has a default of 5. At the point of reporting, we discard strains with a proportion less than a fixed threshold, typically 0.01.
- **Parameters.** In practice, we set the parameters *c* = 100 (Equation (5)); *η* = 0, *σ*^2^ = 3 and *s* = 40 (Sections 2.2.2 and 2.3.1). We set the read error rate as 0.01 and the rate of mis-copying as 0.01.
- **Recombination rate and scaling.** We assume a uniform recombination map, where the genetic distance between loci *i* and *i* + 1 is computed by *ψ_i_* = *D_i_ / d_m_* where *D_i_* denotes the physical distance between loci *i* and *i* + 1 in nucleotides and *d_m_* denotes the average recombination rate in Morgans bp^−1^. We use the recombination rate for *P falciparum* of 15,000 base pairs per centiMorgan as reported by Miles et al. (2016). The recombination rate is scaled by a factor *G*, which reflects the effective population size, rate of inbreeding and size and relatedness of the reference panel. In practice, we have found that a value of *G* = 20 works well. The scaled genetic distance *Gψ* is used to compute the transition probability of switching from copying reference haplotype a to reference haplotype b (see Supplementary Materials for details).
- **Update without linkage disequilibrium.** For initialising the chain, or if the markers present are very widely spaced, linkage disequilibrium can be ignored, which is equivalent to setting the genetic distance between adjacent loci to be infinitely high. Under these circumstances, the haplotype updates become much simpler and depend only on the population-level allele frequency (PLAF), for example as estimated from the reference panel or provided independently.
- **Reporting.** We aim to provide users with a single point estimate of the haplotypes and their proportions, although the full chain is also available for analysis. To achieve this we report values at the last iteration - i.e. we report a single sample from the posterior. However, to measure robustness, we also typically repeat deconvolution with multiple random starting points and select the chain with the lowest average deviance (after removing the burn-in) to report. The deviance measures the difference in log likelihood between the fitted and saturated models, the latter being inferred by setting the WSAF to that observed. These parameters can be modified by users to achieve a preferred balance between computational speed and confidence. By default, we set the MCMC sampling rate as 5, with the first 50% of samples removed as burn in and 800 samples used for estimation.
- **Reference panel construction.** To infer clonal samples for the reference panel we use the Pf3k (Pf3k, 2016) project data, running the algorithm without LD on all samples and identifying those with a dominant haplotype (proportion ¿ 0.99) as clonal. These clonal samples are grouped by region of sampling to form location-specific reference panels. In addition, we have included a number of reference strains, described in more detail below.

## 3 Validation and Performance

As validation we used a set of *in vitro* mixtures created by Wendler (2015) to simulate mixed infections. DNA was extracted from four laboratory parasite lines: 3D7, Dd2, HB3 and 7G8, experimentally mixed in different proportions (see Table 2; figures in brackets), and submitted to the MalariaGEN pipeline (MalariaGEN, 2008) for Illumina sequencing and genotyping (Manske et al., 2012).

**Table 2.** Experimental validation of the `DEploid` method. Inferred percentages (true values in brackets) of the mixed samples. In some cases `DEploid` identifies two near identical strains due to some erroneously called heterozygous sites. The “+” sign indicates the combined proportion.

| sample | 3D7 | Dd2 | HB3 | 7G8 |
| --- | --- | --- | --- | --- |
| PG0389-C | 88.5 (90) | 11.5 (10) | 0 | 0 |
| PG0390-C | 79.8 (80) | 20.2 (20) | 0 | 0 |
| PG0391-C | 66.1 (67) | 33.9 (33) | 0 | 0 |
| PG0392-C | 31.2 (33) | 68.8 (67) | 0 | 0 |
| PG0393-C | 18.4 (20) | 81.6 (80) | 0 | 0 |
| PG0394-C | 9.1 (10) | 90.1 (90) | 0 | 0 |
| PG0395-C | 0 | 33.6 (33.3) | 35 (33.3) | 31.3 (33.3) |
| PG0396-C | 0 | 25.9 (25) | 26.1 (25) | 48 (50) |
| PG0397-C | 0 | 14.7 (14.3) | 15.3 (14.3) | 69.9 (71.4) |
| PG0398-C | 0 | 0 | 45.1+54.9 (100) | 0 |
| PG0399-C | 0 | 0 | 56.7+40.9 (99) | 2.4 (1) |
| PG0400-C | 0 | 0 | 39.5+57.5 (95) | 3 (5) |
| PG0401-C | 0 | 0 | 33.3+56.7 (90) | 10 (10) |
| PG0402-C | 0 | 0 | 85.2 (85) | 14.8 (15) |
| PG0403-C | 0 | 0 | 80.1 (80) | 19.3 (20) |
| PG0404-C | 0 | 0 | 75.4 (75) | 24.6 (25) |
| PG0405-C | 0 | 0 | 70.6 (70) | 29.4 (30) |
| PG0406-C | 0 | 0 | 61 (60) | 39 (40) |
| PG0407-C | 0 | 0 | 50.5 (50) | 49.5 (50) |
| PG0408-C | 0 | 0 | 40.1 (40) | 59.2 (60) |
| PG0409-C | 0 | 0 | 30.1 (30) | 69.1 (70) |
| PG0410-C | 0 | 0 | 25.9 (25) | 73.4 (75) |
| PG0411-C | 0 | 0 | 21.4 (20) | 78.5 (80) |
| PG0412-C | 0 | 0 | 15.2 (15) | 84.8 (85) |
| PG0413-C | 0 | 0 | 3.8 (5) | 96.2 (95) |
| PG0414-C | 0 | 0 | 0 (1) | 29.9+70.1 (99) |
| PG0415-C | 0 | 0 | 0 | 30.0+70.0 (100) |

This data set only contains two unmixed samples, which is insufficient for constructing a reference panel. Moreover, the *P. falciparum* genetic crosses project (Miles et al., 2016) found that due to sequencing error, mapping error and variation among variant calling methods, genotype calls vary at the same locus for the same strain of P *falciparum.* To create a baseline reference haplotype for each strain we therefore considered multiple samples that contains the same parasite strains.

### Inferring haplotypes for Dd2 strain

Since 3D7 is the reference strain, we assume that strain Dd2 is the only source of ‘ALT’ reads in samples PG0389-C to PG0394-C. Assuming markers are independent from each other, let *y* be the read count for ‘ALT’ allele and *x* be the total read count weighted by the Dd2 mixing proportion (see Table 2 in brackets), we use a regression model (*y* = *β*_0_ + *β_1_x*) to infer the Dd2 genotype: 1 if *β*_1_ is significant with *p*-values below 0.001; 0 otherwise.

### Inferring haplotypes for HB3 and 7G8

Similarly, for samples PG0398-C to PG0415-C, we let variables *x*_1_, *x*_2_ be the coverages weighted by the mixing proportions of HB3 and 7G8 respectively; we use a regression model (*y* = *β*_0_ *+ β*_1_*x*_1_ *+ β*_2_*x*_2_) to infer the genotypes of HB3 and 7G8: HB3 is 1 if *β*_2_ is significant with *p*-values below 0.001; 0 otherwise; similarly for 7G8.

### 3.1 Accuracy

#### 3.1.1 Proportions and number of strains

To validate our method we applied `DEploid` to 27 lab-mixed *in vitro* samples. We start by assuming at most three strains present in the mixtures and discard strains with an inferred proportion less than 1%. `DEploid` successfully recovers the proportions with haplotypes of the input (see Table 2). The deviation between our proportion estimates and the truth is at most 2%.

However, we also found that in some cases, `DEploid` fits additional strains. For example, in Table 2, we infer six of the HB3 and 7G8 mixtures as mixing of three. On further inspection, two inferred strains are near identical, but separate 9/10/20189/10/2018d because of a few heterozygous sites with high coverage resulting in high leverage in our model (Supplemental Material Figure S3.3(a)). These sites are likely artefacts arising from duplicated sequence that is absent from the reference strain. Such erroneous markers are not currently inferred by `DEploid`, though this could be included in future versions.

To investigate how the accuracy of haplotype inference is affected by the quality of the reference panel (in terms of having haplotypes close to those present in the samples) we experimented with deconvolving the 27 lab-mixed samples with the following reference panels:

- panel I: five Asian and five African clonal strains from the Pf3k(Pf3k, 2016) resource: PD0498-C, PD0500-C, PD0660-C, PH0047-Cx, PH0064-C, PT0002-CW, PT0007-CW, PT0008-CW, PT0014-CW, PT0018-CW.
- panel II: panel I with the addition of HB3;
- panel III: panel II with the addition of 7G8;
- panel IV: panel III with the addition of Dd2;
- panel V: 3D7, HB3, 7G8 and Dd2 strains (the perfect reference panel for the lab mixtures).
- panel VI: panel I with the addition of six (three each) clonal strains from Asia and Africa: PH0193-C, PH0283-C, PH0305, PT0060-C, PT0146-C and PT0158-C (a typical reference panel for field samples of unknown geographical origin).

In all cases we estimated the number and proportion of strains accurately, for example Figure 2 shows the proportions of strains Dd2/7G8/HB3 as being accurately inferred as approximately 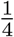, 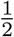, and 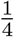.

**Figure 2.**
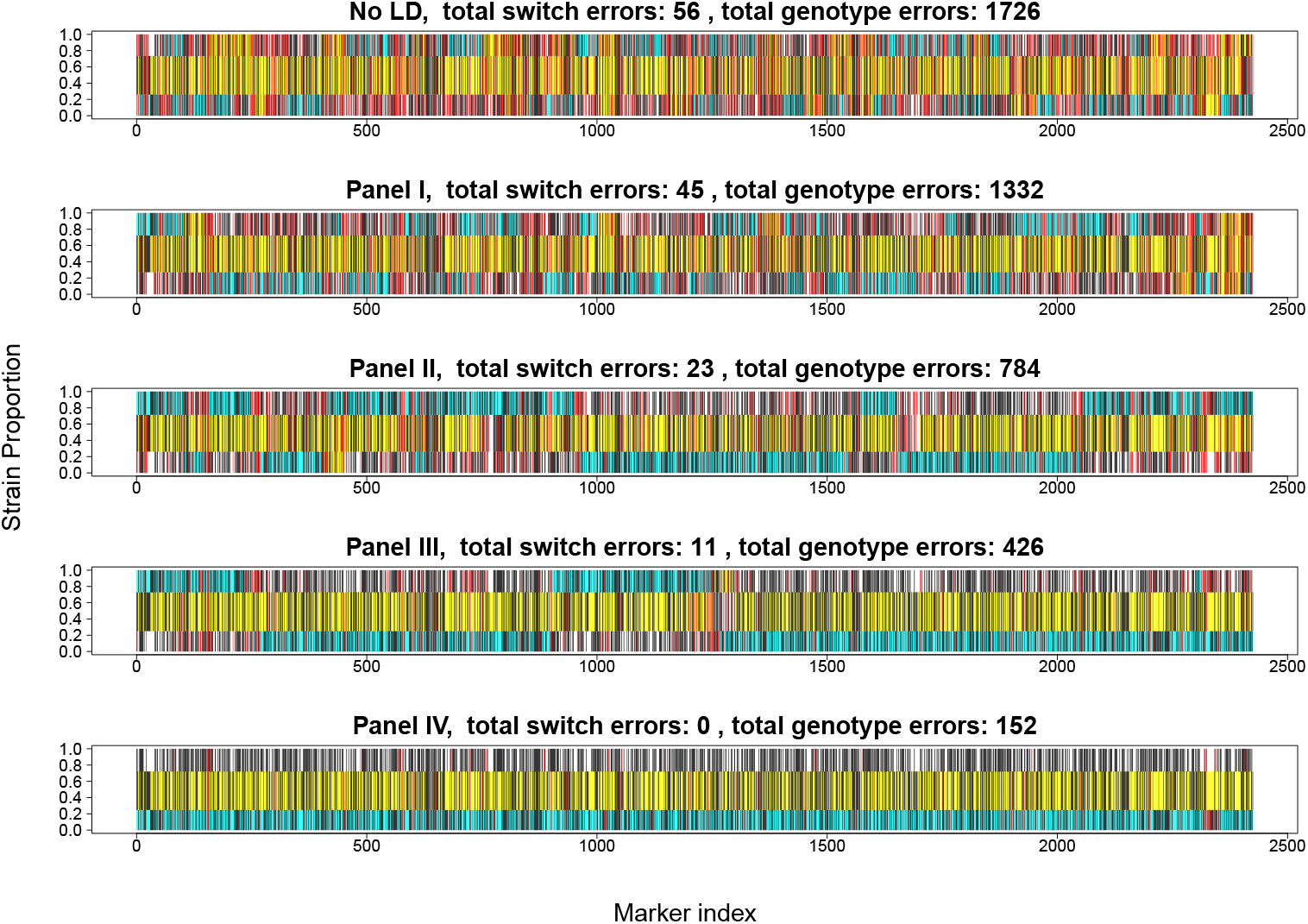
Comparison of true and inferred haplotypes for Chromosome 14 in sample PG0396-C without linkage disequilibrium (top) and using Reference Panels I to IV (from the second to the bottom). Reference Panel V gives results equivalent Panel IV and Panel VI gives results similar to Panel I. Black bars indicate alternative alleles; red bars mark wrongly inferred positions. The yellow, cyan and white background label the haplotype segments from strains 7G8, HB3 and Dd2 respectively. The switch errors are obtained by counting the changes of a strain segment mapped to reference strains; the genotype errors are the discordance between the strain and the mapped reference segments.

#### 3.1.2 Haplotypes

Our accuracy assessment for inferred haplotypes takes into account both switch errors and genotype discordance, which reflects recombination and miscopying events. To understand how the inferred haplotypes relates to those present we split haplotypes into sets of 50 consecutive variants and assigned them to the reference strains through maximal identity. Switches occur when adjacent segments of inferred haplotypes are closest to different reference strains. Genotyping errors occur when a subset of sites within the segment differ from the closest reference strain. Example deconvolutions are shown in Figure 2 and an overview of all experiments is shown in Figure 3. From our assessment of haplotype inference, we conclude:

- The inference of relative proportions does not seem to be affected by the use of linkage disequilibrium information from the reference panel or its closeness to the samples being analysed (Figure 2).
- The accuracy of haplotype inference is, however, dependent on having an appropriate reference panel in terms of relatedness to the samples being analysed (Figure 2).
- The strain proportion affects haplotype inference (see Figure 3). Our method infers strains with proportions over approximately 20% with high accuracy, but struggles with minor strains due to insufficient data, in particular at sites when the minor strain carries the alternative allele and the dominant strain carries the reference allele (see Figure 3).

**Figure 3.**
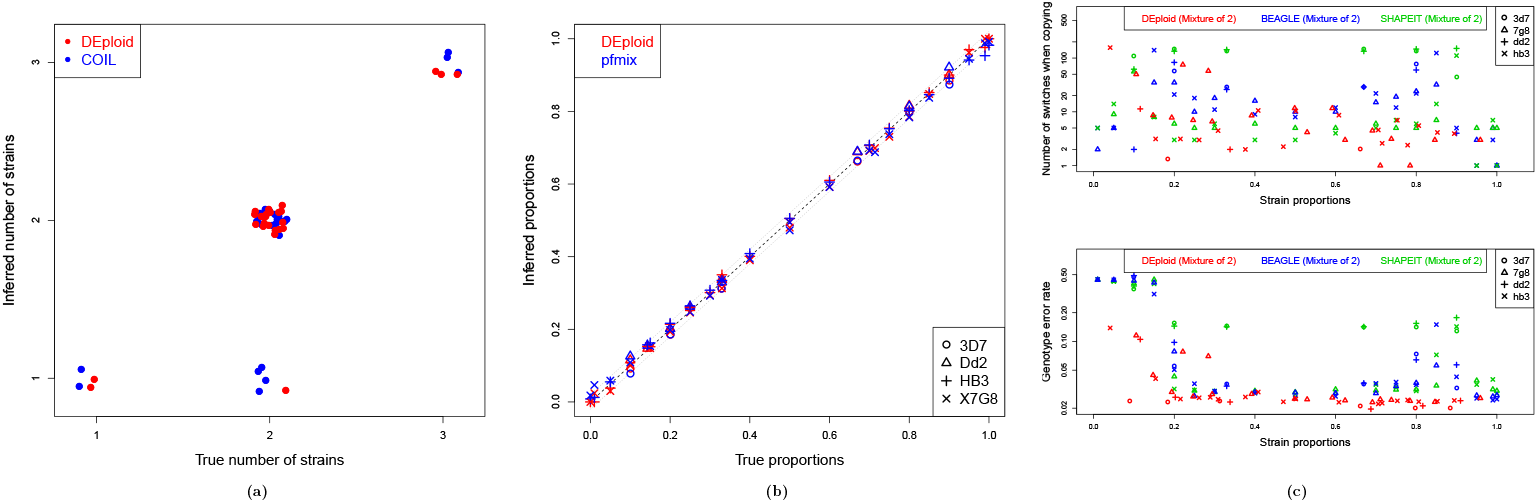
Comparison of `DEploid` and existing tools (`COIL, pfmix`, `BEAGLE`, and `SHAPEIT`). (a) Estimates for the number of strains present in each mixed infection (artificially mixed in the lab) as given by `COIL` and `DEploid`. (b) Comparison of the proportion estimates of each strain as given by `pfmix` and `DEploid`. (c) Relationship between strain proportions and haplotype inference accuracy in the experimental validation for `DEploid` and `BEAGLE/SHAPEIT` (only mixtures of two strains). We use reference panel V to deconvolute all 27 samples. Each point represents a deconvoluted haplotype with 18,570 sites. Point shape refers to strain and colour indicates the method applied. Top panel shows switch error rate whereas the bottom panel indicates genotyping error rate.

### 3.2 Comparison to existing methods

A mixed infection can be completely described by the number of co-existing strains, their relative proportions, and their associated haplotypes. Existing methods for characterizing mixed infections are limited to providing a summary statistic of relative inbreeding (F_ws_,Manske et al. (2012)), inferring the number of strains (`COIL`), or simultaneously inferring the number of strains and their proportions (pfmix, O’Brien et al. (2015)). `DEploid` is the only method that can also estimate haplotypes although it can be argued that conventional tools for phasing diploid organisms (`BEAGLE, SHAPEIT`) could be used to deconvolute mixtures of two strains.

In this section, we use the same dataset (27 samples) to compare `DEploid` with all the inferential methods mentioned above (see Supplementary Material for details). Our method shows robust inference on the number of strains when relative proportions are above 1%. `DEploid` correctly infers the number of strains in 26 out of 27 samples.

In comparison, `COIL` correctly infers the number of strains in 23 samples. We notice that `COIL` struggles to identify strains whose relative proportions is below 5% (Figure 3(a)). Regarding the inference of relative proportions, `DEploid` and pfmix produce similar results with minor differences (Figure 3(b)), with pfmix deviating from the truth at most 3.6%, and `DEploid` exhibiting a maximum error of 2%.

We also experimented with `BEAGLE` and `SHAPEIT` for deconvolving haplotypes in mixtures of two strains. Both methods worked well for balanced mixtures (i.e. with proportions between 40% and 60%) as they mimic a diploid sample. However, as strain proportions became more unbalanced, accuracy degraded and both methods wrongly inferred heterozygous sites as homozygous, introducing a bias towards inferring the haplotypes of dominant strains. We observed that strains with a relative proportion below 20% were always masked out by the dominant strain (Figure 3(c)).

### 3.3 Run-time

The complexity of our program is *𝒪* (*lm*^2^) (see Figure 4), where *m* and *l* are the number of reference strains and sites respectively. In practice, we recommend dividing samples into distinct geographical regions to perform deconvolution, using the ten most different local clonal strains as as reference panel. The run time for deconvolution a field sample range between 1 and 6 hours, depending on the number variants in a sample: For example, it takes 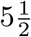 hours to process sample QG0182-C over 372,884 sites. We give worked examples of deconvolving mixed infections from *in vitro* samples in the Supplementary Material.

**Figure 4.**
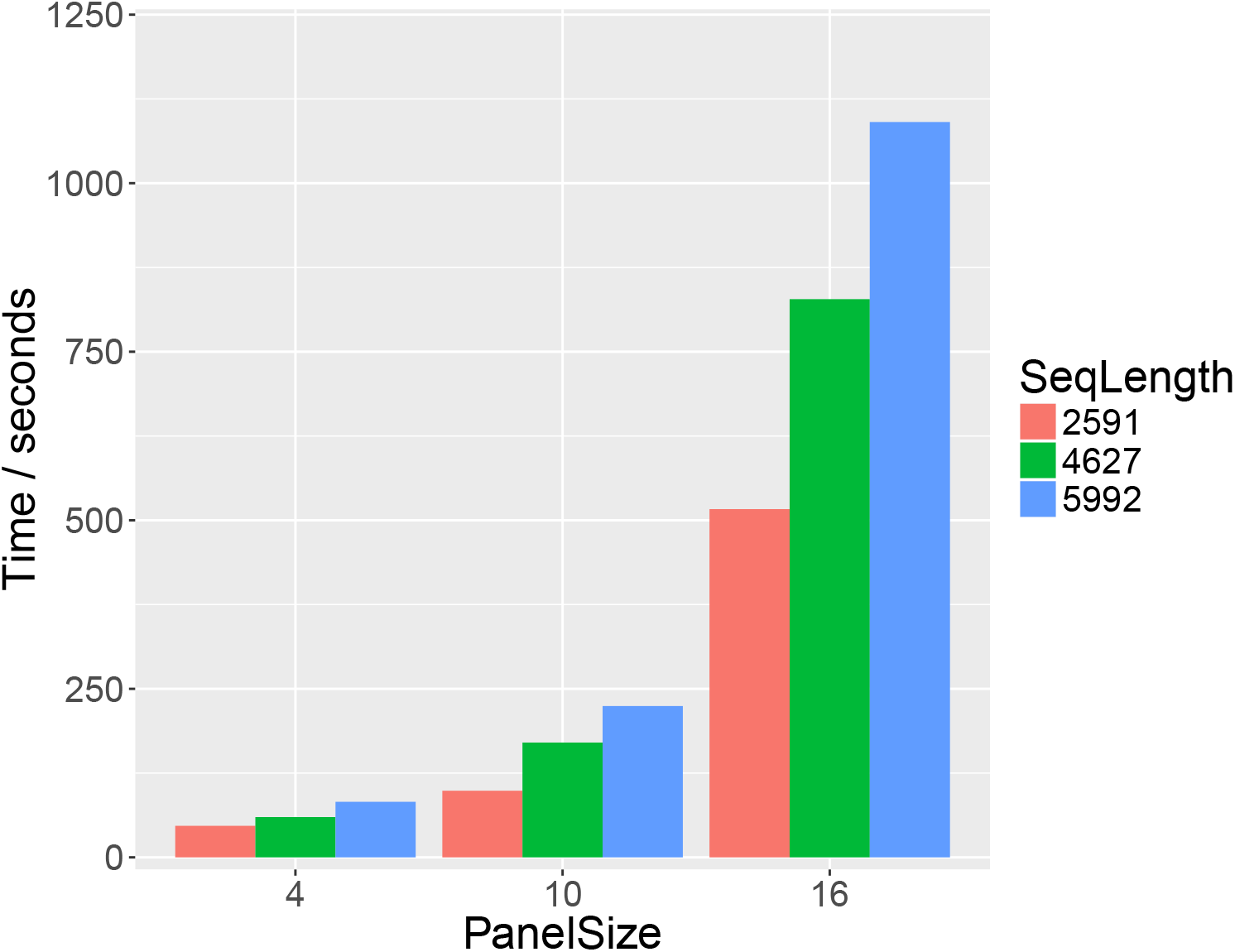
Run-time and scaling. CPU time (seconds) for deconvolving chromosomes 12, 13 and 14 of sample PG0412-C with reference panels I, V and VI (size 4, 10 and 16 reference haplotypes respectively). The run-time is approximately linear with respect to the number of sites and shows the expected quadratic trend against the number of reference strains.

## 4 Discussion

The program `DEploid` and its analysis pipeline has been originally developed for *P. falciparum* studies. Nonetheless, with minor parameter changes, `DEploid` can be used for deconvolution of any other data set with a mixture of samples from a single species, for example on data from *Plasmodium vivax* (Pearson et al., 2016) or bacterial and viral pathogens.

There are several limitation of the current implementation, the greatest of which is the quadratic scaling with reference panel size. In practice, current approaches to related problems such as haplotype phasing (Delaneau et al., 2012) or inference from low-coverage sequencing experiments (Davis, 2016) typically aim to select a few candidate haplotypes (which might be a mosaic) from a reference panel. Alternatively, the reference panel data can itself be approximated, for example through graphical structures, as in BEAGLE (Browning and Browning, 2007), or represented through structures that enable efficient computation (Lunter, 2016). Such extensions will be pursued in future work. Similarly, the observation that a small number of heterozygous sites can lead to inferring the presence of closely related strains should be addressed. Although, in some cases, such sites will reflect *in vivo* evolution, typically most will be erroneous calls and should be identified automatically and excluded.

## Acknowledgements

We thank the Pf3k consortium for valuable insights, in particular, suggestions from Roberto Amato, John O’Brien, Richard Pearson, Jerome Kelleher and Jason Wendler for providing the data of artificial samples. We thank Zam Iqbal for suggesting the name DEploid.

## Funding

Funded by the Wellcome Trust grant [100956/Z/13/Z] to GM.

*Conflict of Interest:* none declared.

